# Root surface treatment for delayed replantation of avulsed teeth in animal models: a systematic review

**DOI:** 10.1101/2024.03.24.586399

**Authors:** Stephanie Díaz Huaman, Marina Moscardini Vilela, Paulo Nelson-Filho, Andiara De Rossi, Alexandra Mussolino de Queiroz, Francisco Wanderley Garcia Paula-Silva

## Abstract

The aim of this study was to perform a systematic review of the *in vivo* effectiveness of different types of root surface treatment materials used in delayed replanted teeth following tooth avulsion in animal models. A systematic review was conducted according to the PRISMA statement. Two reviewers performed a database search for studies published between January 1966 and April 2019 which were indexed in the PubMed, Scopus, and Bireme databases. Studies performed *in vivo*, in animal models with an avulsion/delayed replantation design (≥ 20 min of extra oral dry time) that evaluated the use of different materials for root surface treatment were included. The assessment for risk of bias was performed following recommendations included in Cochrane handbook for systematic reviews of interventions. We found 21 types of materials used for root surface treatment alone and 29 materials used with associations. Stannous fluoride, sodium fluoride, citric acid, doxycycline, Emdogain, alendronate, minocycline, Odanacatib, MFR buffer, recombinant human bone morphogenetic protein, gallium nitrate, acidulated phosphate fluoride, vitamin C, propolis, zoledronic acid, diode laser, indomethacin, fibrin sealant, adipose-tissue derived stem cells treatment and basic fibroblast growth gel. After Grading of Recommendations Assessment, Development, and Evaluation, four studies were scored as low quality of evidence, fifteen studies with moderate quality and six with high quality of evidence. Meta-analysis was not performed due to heterogeneity among studies and materials used for root surface treatment and therefore it was not possible to ascertain which material or protocol present better efficacy when used as root surface treatment material.

## INTRODUCTION

Dental trauma is one of the most common urgencies treated in oral health care centers worldwide. ^1,2^ Among them, avulsion is the most harmful, since the displacement of a dental piece out of its socket due to traumatic forces, causes more serious and irreversible damage to periodontal tissues, needing to be managed as soon as it happens. When a dental piece is avulsed, periodontal structure become compromised, causing disruption of the periodontal ligament (PDL) and neurovascular bundle, and exposing periodontal tissue to environmental contamination.^3^

The aim of replantation is to guarantee the vitality of periodontal ligament and dental pulp; however, its success relies in different associated factors that influence the treatment outcome. When the tooth remains a long period out of socket (longer than 60 minutes), periodontal ligament cells and pulp necrosis may occur, tooth resorption and an eventual tooth loss.^4–6^ Despite the impaired outcomes, delayed replantation has been contemplated as a treatment option by the IADT in complicated case scenarios, mainly because of aesthetics reason.^3,6,7^ Delayed replantation has been reported to lead to mineralized tissue resorption process caused by inflammation and tooth loss later on. In light of that, several materials and protocols have been tested such as root surface treatment aiming to support a suitable response of the periodontal tissue aiming to reduce inflammatory resorption and improve the prognosis.

Root surface treatment materials may be an aid in preventing or delaying the root resorption process when delayed replantation is performed. Several materials have been proposed to be applied as root surface treatment, such as fluorides, citric acid, bisphosphonates, enamel matrix derivative (Emdogain), antibiotics, laser photomodulation therapy, propolis, vitamin C, non-steroidal anti-inflammatory drugs (NSAIDs), calcium hydroxide, gallium nitrate, stem cell therapy, among others. However, there is no consensus in which one the best material would be to prevent sequels. Therefore, the aim of this study was to perform a systematic review of the *in vivo* effectiveness of different types of root surface treatment materials used in delayed replantation in animal models.

## MATERIAL AND METHODS

### Protocol

This systematic review was conducted in accordance with the recommendations of the Preferred Reporting Items for Systematic Reviews and Meta-analysis (PRISMA) statement.

### Focused question

What is the best root surface treatment material or technique to be used in delayed replantation following tooth avulsion?

### Literature search and eligibility criteria

The articles included in this study were obtained in the databases MEDLINE (1966 – December 2019). The search strategy was based on the following Medical Subject Heading terms (MeSH), Text words [tw] and combination strategies: “replantation” [MeSH term] OR “reimplantation” [tw] AND “teeth” [MeSH term] OR “tooth” [MeSH term] AND “avulsion”.

For assessing the studies after MeSH terms search, articles were chosen initially by title, then by abstract and finally, full text article was read to evaluate its relevance in this review (Figure 1). The search was performed by two examiners (SDH and MMV) and in case of disagreement about articles’ importance, a third examiner made the final decision (FWGPS). The articles were chosen based on inclusion criteria such as papers fully written in English, studies performed *in vivo* in animal model with an avulsion/delayed replantation design (≥ 20 min of extra oral dry time) that evaluated the use of different materials for root surface treatment in not endodontically or periodontally-compromised teeth, presence of a control group that represented teeth with late replantation without root surface treatment.

**Figure 1.** Flow diagram illustrating the selection process of studies regarding root surface treatment and delayed tooth replantation.

### Data extraction

The assessment of heterogeneity was performed by two examiners (SDH and MMV) including study design features such as following: animal species, root formation stage, control group, type of teeth, extra oral dry time, removal of necrotic periodontal ligament, material used for root surface treatment, material used for root canal treatment, splinting regimen, follow-up period and outcome measurement. Both authors extracted all the relevant data from the selected studies and organized them into tables. In order to appraise the effectiveness of the material employed for root surface treatment of delayed replanted teeth, data referring periodontal ligament status, presence of inflammation and presence of resorption was extracted.

### Quality assessment and level of evidence

The assessment for risk of bias was performed by two examiners (SDH and FWGPS) following recommendations included in Cochrane handbook for systematic reviews of interventions (Table 1). Included studies were assessed considering the following criteria: randomization, completeness follow-up, balanced experimental group and outcome reporting. The studies were rated as having high, moderate, low, and very low quality of evidence according to the sum of scores based on the GRADE (Table 2) (Grading of Recommendations Assessment, Development, and Evaluation) system.^8^

**Table 1.** Protocol for qualitative scoring of selected studies.

| Subgroups | Item | Component | Classification | Score | Definition |
| --- | --- | --- | --- | --- | --- |
| Clinical aspects | 1. Treatment guidelines for permanent teeth | Root canal treatment | Yes<br>No | 1<br>0 | Studies that performed root canal treatment |
|  |  | Administer systemic antibiotics | Yes<br>No | 1<br>0 | Studies that administered systemic antibiotics |
|  |  | Apply a flexible splint | Yes, or none for rats/mice<br>No | 1<br>0 | Studies that performed splinting |
| Methodological aspects | 2. Sample | Animal model | Monkeys | 2 | Studies that used rodent were scored lower due to the evolutionary distance to humans |
|  |  |  | Dogs | 1 |  |
|  |  |  | Rats | 0 |  |
|  | 3. Interventions | Extra-alveolar time | ≥ 60 minutes | 2 | Studies that described extra-alveolar time and immersion period in media prior to replantation |
|  |  |  | < 60 minutes | 1 |  |
|  |  |  | Unclear | 0 |  |
|  |  | Root formation stage | Clear | 1 | Studies that described the root formation stage (open and closed apex) |
|  |  |  | Unclear | 0 |  |
|  |  | Removal of PDL | Removed | 1 | Studies that performed removal of PDL |
|  |  |  | Not removed/<br>unclear | 0 |  |
|  | 4. Outcomes | Completely defined | Clear<br>Unclear | 1<br>0 | Studies that clearly described the outcome(s) |
|  | 5. Control groups | Definition of control group | Delayed replantation without treatment<br>Immediate replantation without treatment<br>Unclear/not reported/<br>inappropriate control group | 2<br>1<br>0 | Studies that did not include a control group, did not clearly describe the control group, or did not have an appropriate control group were scored lower |
|  | 6. Statistical methods | Statistical methods used | Adequate<br>Partial<br>Inadequate/<br>Inexistent | 2<br>1<br>0 | The statistical analysis was satisfactory to evaluate the data |
|  | 7. Results | Data presentation | Adequate<br>Partial<br>Inadequate | 2<br>1<br>0 | The description of the results was clear and data on different root surface treatment material was fully presented |

**Table 2.** Criteria used for grading quality of evidence of the included studies.

| **GRADE range** | **GRADE quality of evidence** | **Interpretation** |
| --- | --- | --- |
| 13 – 16 | High | Confident that the true effect lies close to the estimated effect. |
| 9 – 12 | Moderate | The true effect is likely to be close to the estimated effect, but there is a possibility that it is substantially different. |
| 5 – 8 | Low | The true effect may be substantially different from the estimated effect. |
| 1 – 4 | Very low | The true effect is likely to be substantially different from the estimated effect. |

Note: Based on GRADE (Grading of Recommendations Assessment, Development, and Evaluation) guidelines.<sup>8</sup>

A meta-analysis was not conducted because the included studies did not have data similar enough to be pooled.

## RESULTS

The databases search resulted in 954 articles and forty-two remained for final full text evaluation (Figure 1). Articles with a title not related to the topic and unclear abstracts were not included. Although, all manuscripts dealt with avulsion and replantation, seventeen articles were discarded due to a different extra oral time, different experimental procedures, evaluation of storage medium instead of root surface treatment and administration of systemic medication that could affect the outcome. Finally, 25 articles were included into this review.

After GRADE evaluation, we scored four studies as low quality of evidence, fifteen studies with moderate quality and six with high quality of evidence (Table 3). Between the studies graded as low quality, we found that treatment protocol used to perform root surface treatment had difference to the IADT Guidelines recommendation, mainly among earlier studies where much information and biological processes were unknown at the time.

**Table 3.** Quality assessment of selected studies.

|  | 1. Treatment guidelines for permanent teeth |  |  | 2. Sample | 3. Interventions |  |  | 4. Outcomes | 5. Control groups | 6. Statistical methods | 7. Results | 8. Quality of evidence | 9. Score |
| --- | --- | --- | --- | --- | --- | --- | --- | --- | --- | --- | --- | --- | --- |
|  | Root canal treatment | Administer systemic antibiotics | Apply flexible splint | Animal model | Extra-alveolar time | Root formation stage | Removal of PDL | Completely defined | Definition of control group | Statistical methods | Data presentation |  |  |
| Bjorvatn & Massler 1971<sup>9</sup> | 0 | 0 | 1 | 1 | 2 | 1 | 0 | 0 | 2 | 0 | 0 | 7 | Low |
| -Klinge et al. 1984<sup>10</sup> | 0 | 0 | 0 | 2 | 1 | 1 | 1 | 1 | 2 | 1 | 1 | 10 | Moderate |
| Bjorvatn et al. 1989<sup>11</sup> | 0 | 0 | 0 | 2 | 1 | 0 | 0 | 1 | 0 | 1 | 2 | 8 | Low |
| Cvek et al. 1990<sup>12</sup> | 0 | 1 | 1 | 3 | 2 | 0 | 0 | 1 | 2 | 1 | 2 | 13 | High |
| Selvig et al. 1992<sup>13</sup> | 0 | 1 | 0 | 2 | 1 | 0 | 0 | 0 | 2 | 1 | 2 | 10 | Moderate |
| Iqbal et al. 2001<sup>14</sup> | 1 | 1 | 0 | 2 | 2 | 1 | 0 | 1 | 2 | 1 | 2 | 13 | High |
| Bryson et al. 2003<sup>15</sup> | 1 | 0 | 0 | 2 | 2 | 1 | 0 | 1 | 2 | 1 | 1 | 11 | Moderate |
| Khin Ma et al. 2003<sup>16</sup> | 1 | 1 | 0 | 3 | 2 | 1 | 0 | 1 | 2 | 1 | 2 | 14 | High |
| Lam e<br>Sae-Lim,<br>2004<br><sup>17</sup> | 1 | 1 | 0 | 3 | 2 | 1 | 1 | 1 | 2 | 1 | 2 | 15 | High |
| Soren<br>sen et<br>al.<br>2004<br><sup>18</sup> | 0 | 1 | 1 | 2 | 1 | 0 | 1 | 1 | 0 | 0 | 1 | 8 | Low |
| Lusto<br>sa-Pereir<br>a et<br>al.<br>2006<br><sup>19</sup> | 1 | 1 | 1 | 1 | 2 | 1 | 1 | 1 | 1 | 0 | 1 | 11 | Mod<br>erate |
| Mori<br>et al.<br>2007<br><sup>20</sup> | 1 | 1 | 1 | 1 | 1 | 1 | 0 | 0 | 0 | 0 | 1 | 7 | Low |
| Poi et<br>al.<br>2007<br><sup>21</sup> | 1 | 1 | 1 | 1 | 2 | 1 | 1 | 1 | 0 | 0 | 1 | 10 | Mod<br>erate |
| Panza<br>rini et<br>al.<br>2008<br><sup>22</sup> | 1 | 1 | 1 | 1 | 2 | 1 | 1 | 0 | 0 | 1 | 1 | 10 | Mod<br>erate |
| Gulin<br>elli et<br>al.<br>2008<br><sup>23</sup> | 1 | 1 | 1 | 1 | 2 | 1 | 1 | 0 | 2 | 1 | 1 | 12 | Mod<br>erate |
| Mori<br>et al.<br>2010<br><sup>24</sup> | 1 | 1 | 1 | 1 | 1 | 1 | 1 | 1 | 0 | 1 | 2 | 11 | Mod<br>erate |
| Carva<br>lho et<br>al.<br>2012<br><sup>25</sup> | 1 | 1 | 1 | 1 | 2 | 1 | 1 | 1 | 2 | 1 | 2 | 14 | High |
| Zanett<br>a-Barbo<br>sa et<br>al.<br>2014<br><sup>26</sup> | 1 | 1 | 0 | 2 | 1 | 0 | 0 | 1 | 2 | 1 | 2 | 11 | Mod<br>erate |
| Panza<br>rini et<br>al.<br>2014<br><sup>27</sup> | 1 | 1 | 1 | 1 | 2 | 1 | 0 | 1 | 2 | 1 | 2 | 13 | High |
| Jabari<br>far et<br>al.<br>2015<br><sup>28</sup> | 0 | 1 | 1 | 2 | 2 | 1 | 0 | 1 | 0 | 1 | 1 | 10 | Mod<br>erate |
| Barbi<br>zam<br>et a.<br>2015<br><sup>29</sup> | 1 | 0 | 1 | 1 | 2 | 1 | 0 | 1 | 2 | 2 | 2 | 13 | High |
| Demir<br>el et<br>al.<br>2016<br><sup>30</sup> | 1 | 1 | 1 | 1 | 2 | 1 | 0 | 1 | 2 | 1 | 2 | 13 | High |
| Carva<br>lho et<br>al.<br>2017<br><sup>31</sup> | 1 | 1 | 1 | 1 | 2 | 1 | 1 | 1 | 2 | 1 | 2 | 14 | High |
| Kwon<br>et al.<br>2018<br><sup>32</sup> | 1 | 1 | 1 | 1 | 2 | 1 | 0 | 1 | 2 | 2 | 2 | 14 | High |

The relevant study design features were organized in Table 4. The features included were animal species studied, root formation stage, control group, intervention characteristics such as: type of teeth – multi-rooted or single rooted, extra oral dry time, method for removal of necrotic periodontal ligament, material used for root surface treatment, material used for endodontic treatment, splinting regimen, follow-up time and outcome measurements.

**Table 4.**
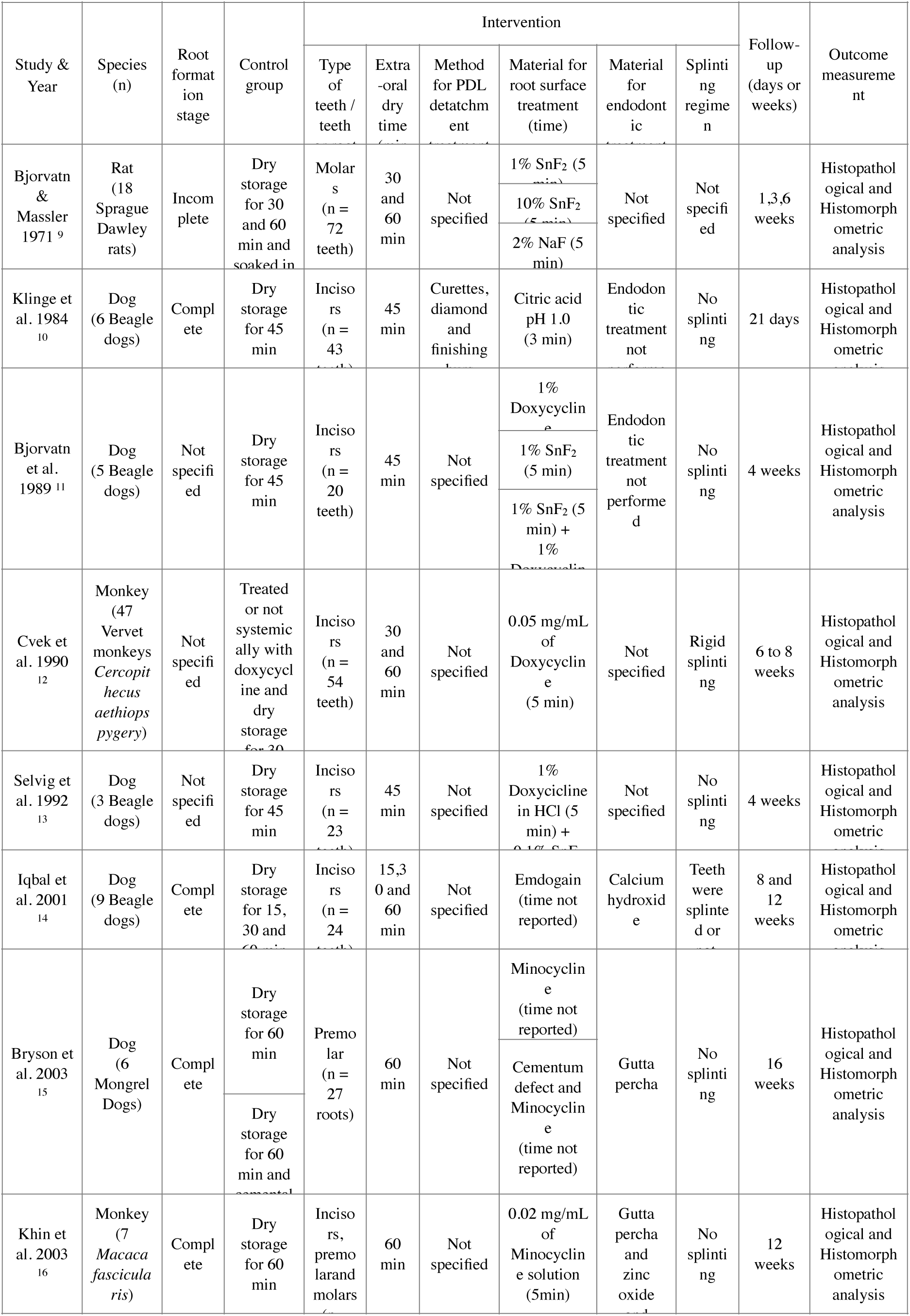
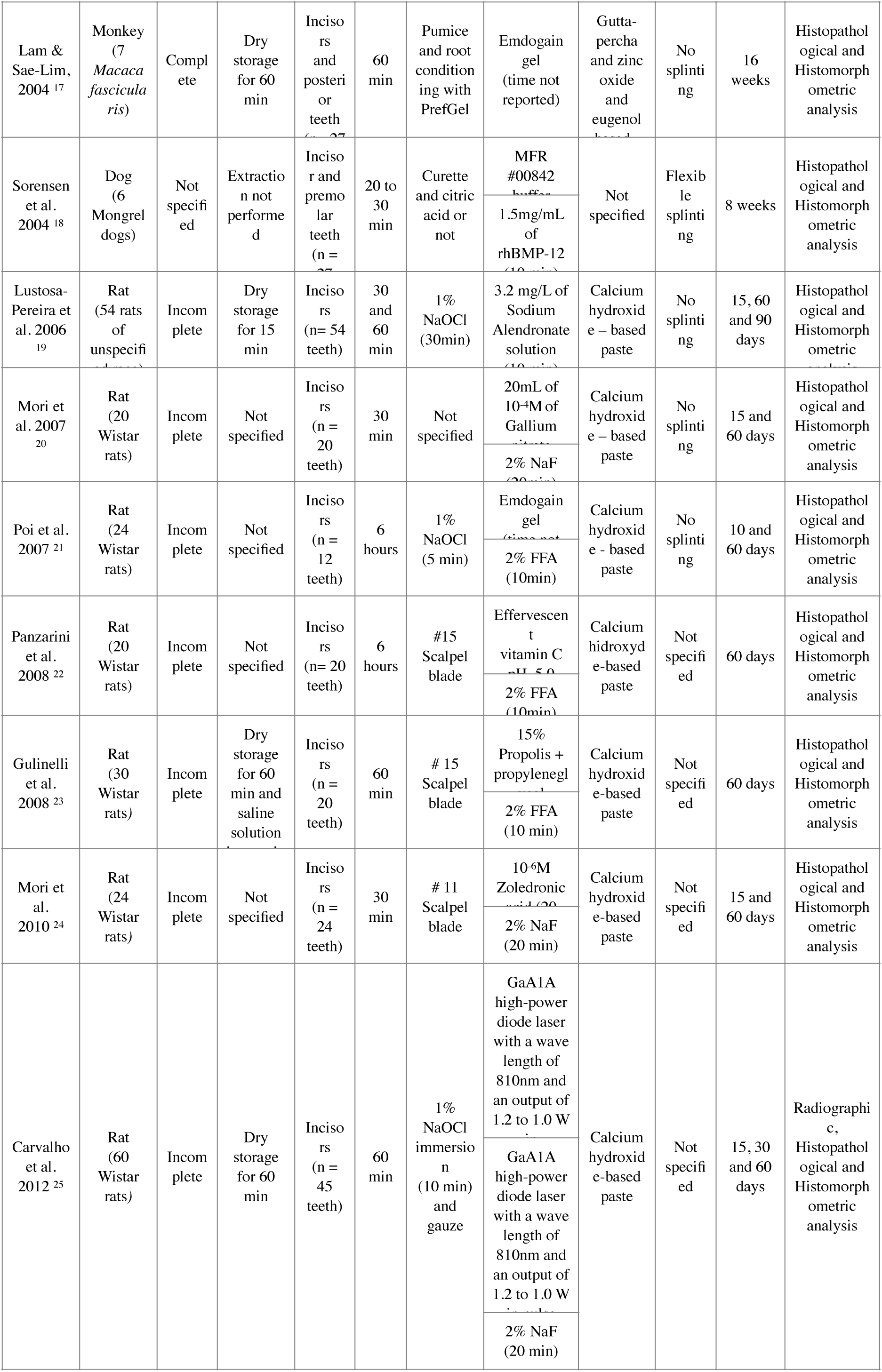
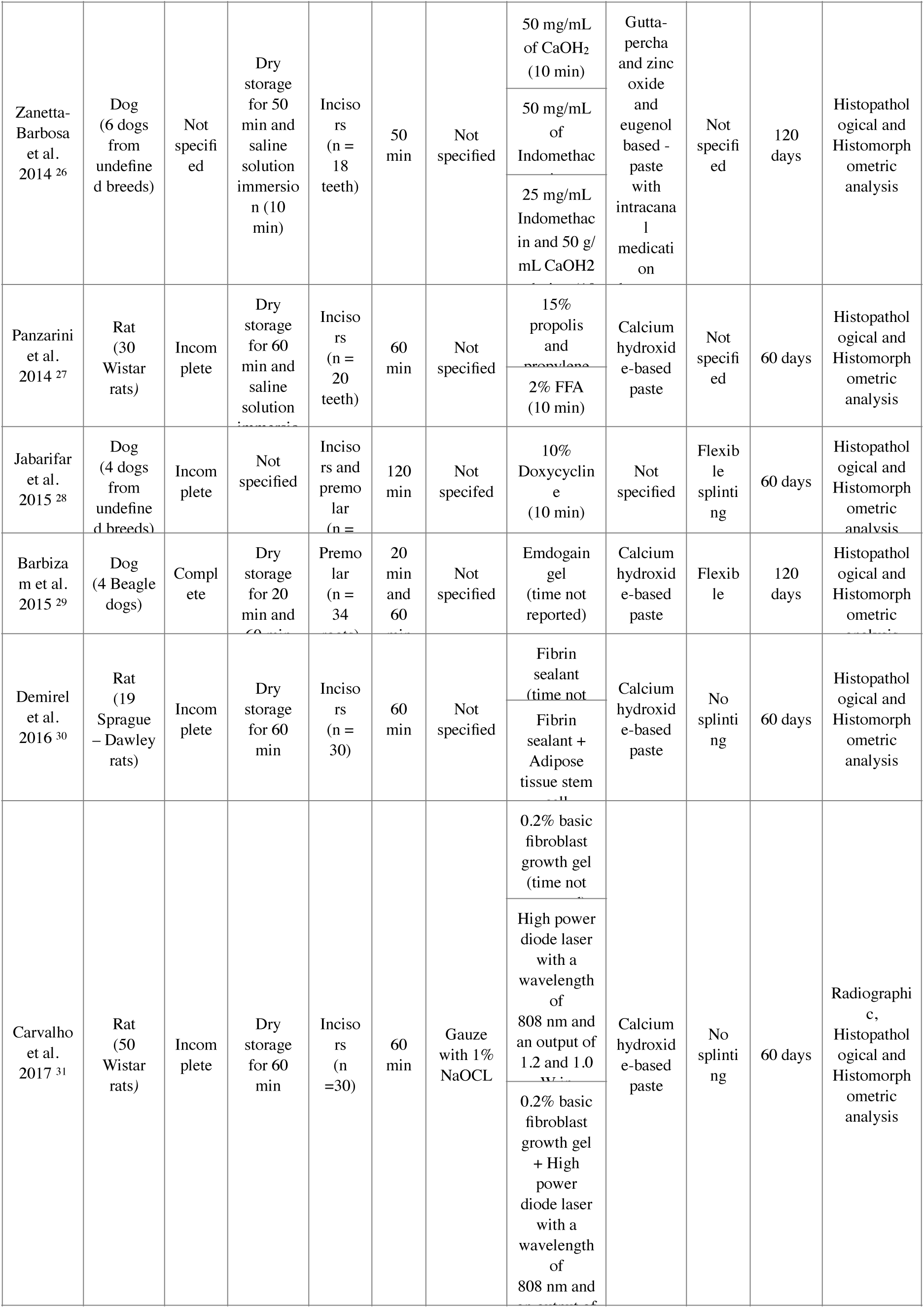
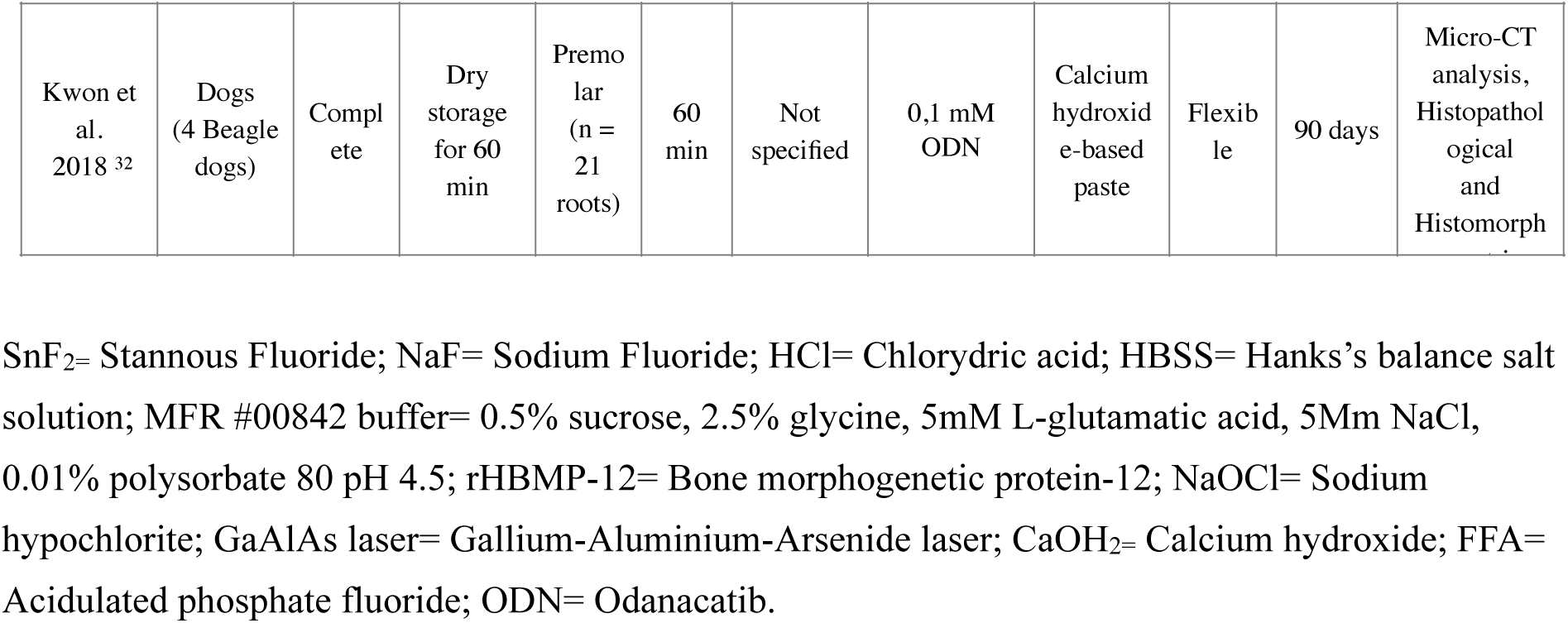
Description of selected studies.

The selected studies were published from 1971 to 2019, in which 11 studies used rats as experimental animals (Sprague Dawley, Wistar or breed not described), 11 used dogs of different breeds (Beagle, Mongrel or undefined breed) and 3 studies used monkeys (Macaca and Vervet monkeys).

The interventions were performed using different types of teeth, predominantly incisors with incomplete root formation. Five studies did not report the root formation stage of the experimental teeth. When molars or premolars were included, tooth section was performed before extraction; nevertheless, it was not always detailed in the article. Before the intervention, teeth were left to dry on a bench, from 15 min to 6 hours. Only in 10 studies were specified the method used for periodontal ligament detachment.

We found 21 types of materials used for root surface treatment alone and 29 materials used with associations. Stannous fluoride (SnF_2_), sodium fluoride (NaF), citric acid, doxycycline, Emdogain, alendronate, minocycline, Odanacatib, MFR buffer (5mM L-glutamic acid, 2.5% glycine, 0.5% sucrose, 0.01% polysorbate 80, pH 4.5; Wyeth Research), recombinant human bone morphogenetic protein (rhBMP), gallium nitrate, acidulated phosphate fluoride (FFA), vitamin C, propolis, zoledronic acid, diode laser, indomethacin, fibrin sealant, adipose-tissue derived stem cells treatment and basic fibroblast growth (bFGF) gel. Doxycycline, FFA 2% and NaF 2% were the materials that were used as experimental groups in most of the studies.

As for results, only 9 studies reported presence of inflammation in the experimental group, meanwhile the information was missing in 18 articles. Ankylosis was reported in 9 studies and only 7 studies reported the periodontal ligament fibers orientation after healing. Ankylosis information was missing in other articles. “Normal Healing” as a feature was used to describe periodontal ligament of the experimental group in 7 studies. “Surface resorption” and “Replacement resorption” were found to be used equally and with the same meaning between included articles. However, its presence was reported only in 8 studies, frequently accompanied by inflammatory resorption.

Statistics analysis differed among the studies, however only one study specified about needing to considerate cluster readjustment and statistics when experimental teeth are included in more than one animal. Some studies mentioned statistics processes performance. Two studies did not performed statistics analysis being graded with a low grade.

Meta-analysis was not performed due to heterogeneity between the studies and materials used for root surface treatment.

## DISCUSSION

In recent years, root surface treatment protocol has been a controversial subject of discussion. Although the use of sodium fluoride 2% (NaF) for coating the root surface before delayed replantation had been previously recommended,^3^ currently there is no evidence to establish NaF as a gold standard between root surface treatment materials. Novel substances that may potentially delay replacement resorption and decrease the possibility of adverse outcomes such as inflammatory resorption had been reported. In need to reach consensus in which material for root surface treatment could improve the treatment success, we performed a systematic review to identify materials that have been applied in an animal model to determine their efficacy.

After searching databases, we found that root formation stage was not specified in 5 studies, 13 included teeth with incomplete root formation and 6 teeth with complete formation. Previous studies have reported root formation stage as an important predictor of the replantation outcome.^33^ Case reports of immature teeth having better outcomes after replantation have been found in current literature, unfortunately, scientific evidence is still lacking.^6,33^ Information about root formation was not found in all studies included in this review.

Control groups used in these studies was diverse and the extra oral time were different in every study. The IADT guidelines had established that up until 60 minutes could be considered immediate replantation. We included studies that had a control group where root surface was exposed to an extraoral environment up to 60 minutes or that had been immediately replanted. However, in order to have a more comparable results, experimental models should have a positive control group where delayed replantation is performed without any surface treatment. Studies with immediate replantation or with a control group with an unclear treatment or no treatment at all, received lower scores.

We also found that root canal treatment was only performed in some studies. Studies that did not performed root canal treatment were held in the 1980’s decade. It could be presumed that at that time, the relevance of pulp tissue contamination due to dental trauma on periodontal healing was not known. However, now is well known that the presence of bacteria in the root canal will prevent repair and sustain the inflammation in the periodontal ligament.^34^

Systemic administration of antibiotics has been also used to improve periodontal ligament healing. Systemic administration of doxycycline was investigated in one study held by Cvek.^12^ Doxycycline is an antibiotic that has been widely studied to be used in the treatment of avulsed teeth, as storage medium in solution and as a surface treatment material. It has been reported that its topical use may present antimicrobial effect enhancing the replantation success rate.^12,13^ However, when used systemically, doxycycline was not been found to improve the outcome after replantation of avulsed teeth.^12^ On the other hand, administration of amoxicillin or tetracycline had a positive influence on the repair process in delayed tooth replantation.^11^ IADT guidelines recommend antibiotics prescription as a measure of prevention when dental trauma occurred in highly contaminated areas and in situations where cleansing of avulsed tooth was not adequate.^3^ In this review, we considered the inclusion of a systemic antibiotic as a favorable feature in the experimental design since its absence may ultimately affect the outcome.^35^

Periodontal ligament removal treatment was not specified in 13 studies. In 11 studies necrotic periodontal ligament was removed with different methods: curettes, finishing burs, citric acid, NaOCl, scalpel blade and gauze. Periodontal ligament detachment treatment with gauze was found to not provide protection from ankylosis or replacement resorption in delayed replanted teeth.^33^ However, IADT recommends the removal of necrotic periodontal ligament since it was found that presence of necrotic tissue induces osteoclastogenesis and larger resorption areas.^34,35^ Studies that did not specified or do not performed the periodontal ligament detachment treatment received lower scores.

Between the root surface treatment materials that were found. Fluoride solutions such as FFA, SnF_2_ and NaF were constantly used as a control group treatment, probably because of IADT guidelines recommendation. Doxicycline and Minocycline are locally applied antibiotics under the premise that systemic antibiotics do not have any beneficial effect on pulpal or periodontal healing.^36,37^ In this review, we found six studies that used antibiotics, two of them associated to a fluoride. Materials such as FGF, BMP-12 and adipose tissue stem cells in association with fibrin sealant were also used as a root surface treatment. These materials were found to stimulate osteogenic differentiation and promote cell proliferation when applied to human periodontal cells *in vitro* and *in vivo.*^38^

Biphosphonates such as alendronate and zoledronic acid were also used in root canal treatment. Drugs that are applied to prevent bone resorption were used considering a similarity between enzymes, markers and cells involved in bone resorption and those involved in root resorption.^20,24^ Photodynamic therapy (PDT) was also presented as a treatment option between the root surface treatment materials. PDT has a well-known set of advantages: an antimicrobial potency, induce cell proliferation, osteogenesis and endothelial cells proliferation.^39–41^

A flexible splint may provide a more favorable outcome where the periodontal ligament fibers reattach following tooth avulsion.^33^ In rodent teeth, splinting after replantation is not strictly recommended since the anatomy of the socket provides a stable position.^34^ We found that 12 studies included in this review did not performed splinting, from these, 11 studies where performed in rats. As IADT recommends in treatment for human teeth, we considered flexible splinting as a mandatory step to achieve healing after replantation, however, 8 studies did not detail if splinting was performed, 1 study in dogs performed rigid splinting and only 4 studies performed flexible splinting. Thus, studies without splinting included were considered of lower quality.

This systematic review has reunited classic and state of art literature about root surface treatment materials having encountered a wide variety of substances that have been tested to prevent mineralized tissue resorption and possibly heal periodontal ligament in delayed replanted teeth. However, even though, six articles were scored as high-quality evidence, due to diverse materials used during experimentation, it was not possible to perform meta-analysis.

In conclusion, we found a high heterogeneity among the studies, not being possible to ascertain which material or protocol present better efficacy when used as root surface treatment material. Nonetheless, this review shed light on methodological issues that should be considered on future research that is deemed necessary in this field.

## ACKNOWLEDGMENTS

This study was financed in part by the Coordenação de Aperfeiçoamento de Pessoal de Nível Superior - Brasil (CAPES) - Finance Code 001 (Fellowships to SDH and MMV).

## REFERENCES

1. Langridge FC, Hufanga SV, Ofanoa MM, Fakakovikaetau T, Percival TM, Grant CC. Child morbidity as described by hospital admissions for primary school aged children in Tonga 2009-2013. N Z Med J. 2017;130(1465):29–43.

2. Mahmoodi B, Rahimi-Ned R, Weusmann J, Azaripour A, Walter C, Willerhausen B. Traumatic dental injuries in a university hospital: a four-year retrospective study. BMC Oral Health. 2015; 15: 139

3. Andersson L, Andreasen JO, Day P, Heithersay G, Trope M, DiAngelis AJ, et al. Guidelines for the Management of Traumatic Dental Injuries: 2. Avulsion of Permanent Teeth. Pediatr Dent. 2017;39(6):412–9.

4. Andreasen JO, Andreasen FM. Avulsions. In: Andreasen JO, Andreasen FM, Andersson L, editors. Textbook and color atlas of traumatic injuries to the teeth, 4th edn. Oxford, UK: Wiley-Blackwell; 2007. p. 444–88.

5. Soder PO, Otteskog P, Andreasen JO, Modeer T. Effect of drying on viability of periodontal membrane. Scand J Dent Res. 1977; 85:164–8.

6. Hupp JG, Mesaros SV, Aukhil I, Trope M. Periodontal ligament vitality and histologic healing of teeth stored for extended periods before transplantation. Endod Dent Traumatol. 1998; 14: 79–83.

7. Kinirons MJ, Gregg TA, Welbury RR, Cole BOI. Variations in the presenting and treatment features in reimplanted permanent incisors in children and their effect on the prevalence of root resorption. Br Dent J. 2000; 189(5): 263–266.

8. Guyatt G, Oxman AD, Akl EA, Kunz R, Vist G, Brozek J, Norris S, Falck-Ytter Y, Glasziou P, DeBeer H, Jaeschke R, Rind D, Meerpohl J, Dahm P, Schünemann HJ. GRADE guidelines: 1. Introduction-GRADE evidence profiles and summary of findings tables. J Clin Epidemiol. 2011;64(4):383–94.

9. Bjorvatn K, Massler M. Effect of fluorides on root resorption in replanted rat molars. Acta Odontol Scand. 1971; 29: 17–29.

10. Klinge B, Nilvéus R, Selvig KA. The effect of citric acid on repair after delayed tooth replantation in dogs. Act Odontol Scand. 1984; 42(6): 351–359.

11. Bjorvatn K, Selvig KA, Klinge B: Effect of tetracycline and SnF2 on root resorption in replanted incisors in dogs. Scand J Dent Res. 1989; 97: 477–82.

12. Cvek M, Cleaton-Jones P, Austin J, Kling M, Lownie J, Fatti P. Effect of topical application of doxycycline on pulp revascularization and periodontal healing in reimplanted monkey incisors. Endod Dent Traumatol. 1990; 6: 170–176.

13. Selvig KA, Bjorvatn K, Bogle GC, Wikesjo UME: Effect of stannous fluoride and tetracycline on periodontal repair after delayed tooth replantation in dogs. Scand Dent Res. 1992; 100: 200–3

14. Iqbal MK, Bamaas NS. Effect of enamel matrix derivative (EMDOGAIN®) upon periodontal healing after replantation of permanent incisors in Beagle dogs. Dent Traumatol. 2001; 17: 36–45.

15. Bryson EC, Levin L, Banchs F, Trope M. Effect of minocycline on healing of replanted dog teeth after extended dry times. Dent Traumatol. 2003;19: 90–95.

16. Khin Ma M, Sae-Lim V. The effect of topical minocycline on replacement resorption of replanted monkey’s teeth. Dent Traumatol. 2003; 19: 96–102.

17. Lam K, Sae-Lim V. The effect of Emdogain gel on periodontal healing in replanted monkey’s teeth. Oral Surg Oral Med Oral Pathol Oral Radiol Endod. 2004; 97:100–107.

18. Sorensen RG, Polimeni G, Kinoshita A, Wozney JM, Wikesjö UME: Effect of recombinant human bone morphogenetic protein-12 (rhBMP-12) on regeneration of periodontal attachment following tooth replantation in dogs. A pilot study. J Clin Periodontol. 2004; 31: 654–661.

19. Lustosa-Pereira A, Garcia RB, de Moraes IG, Bernardineli N, Bramante CM, Bortoluzzi EA. Evaluation of the topical effect of alendronate on the root surface of extracted and replanted teeth. Microscopic analysis on rats’ teeth. Dent Traumatol. 2006; 22: 30–35.

20. Mori GG, Moraes IG, Garcia RB, Borro LCB, Purificação BR. Microscopic Investigation of the gallium nitrate for root surface treatment in rat teeth submitted to delayed replantation. Braz Dent J. 2007; 18(3): 198–201.

21. Poi WR, Carvalho RM, Panzarini SR, Sonoda CK, Manfrin TM, Rodrigues TS. Influence of enamel matrix derivative (Emdogain) and sodium fluoride on the healing process in delayed tooth replantation: histologic and histometric analysis in rats. Dent Traumatol. 2007;23(1):35–41.

22. Panzarini SR, Poi WRM Pedrini D, Sonoda CK, Brandini DA, Luvizuto ER. Root surface treatment with vitamin C in tooth replantation: microscopic study in rats. Minerva Stomatol. 2008; 57:423–428.

23. Gulinelli JL, Panzarini SR, Fattah CMRS, Poi WR, Sonoda CK, Negri MR, Saito CTMH. Effect of root surface treatment with propolis and fluoride in delayed tooth replantation in rats. Dent Traumatol. 2008; 24: 651–657.

24. Mori GG, Janjacomo DMM, Nunes DC, Castilho LR. Effect of zoledronic acid used in the root surface treatment of late replanted teeth: a study in rats. Braz Dent J. 2010; 21(5): 452–457.

25. Carvalho ES, Costa FTS, Campos MS, Ambinder AL, Neves ACC, Habitante SM, Lage-Marques JL, Raldi DP. Root surface treatment using diode laser in delayed tooth replantation: radiographic and histomorphometric analyses in rats. Dent Traumatol. 2012; 28: 429–436.

26. Zanetta-Barbosa D, Moura CCG, Machado JR, Crema VO, Lima CAP, Carvalho ACP. Effect of indomethacin on surface treatment and intracanal dressing of replanted teeth in dogs. J Oral Maxillofac Surg. 2014; 72:127.

27. Panzarini SR, Nonato CC, Gulinelli JL, Poi WR, Sonoda CK, Sato CTMH, Marão HF. Effect of the treatment of root surface-adhere necrotic periodontal ligament with propolis or fluoride in delayed rat tooth replantation. Clin Oral Invest. 2014; 18: 1329–1333.

28. Jabarifar E, Khalighinejad N, Khademi AA, Razavi SM, Birjandi N, Badrian H, Ansari G. Histologic evaluation of apical pulp of immature apex following extraction, surface treatment, and replantation in different storage media in dogs. Dent Traumatol. 2015; 31: 118–124.

29. Barbizam JVB, Massarwa R, Silva LAB, Silva RAB, Nelson-Filho P, Consolaro A, Cohenca N. Histopathological evaluation of the effects of variable extraoral dry times and enamel matrix proteins (enamel matrix derivatives) application on replanted dogs’ teeth. Dent Traumatol. 2015; 31: 29–34.

30. Demirel S, Yalvac ME, Tapsin S, Akyuz S, Ak E, Cetinel S, Yarat A, Sahin F. Tooth replantation with adipose tissue stem cells and fibrin sealant: microscopic analysis of rat’s teeth. Springerplus. 2016; 5: 656.

31. Carvalho ES, Rosa RH, Pereira FM, Anbinder AL, Mello I, Habitante SM, Raldi DP. Effects of diode laser irradiation and fibroblast growth fator on periodontal healing of replanted teeth after extended extra-oral dry time. Dent Traumatol. 2017; 33: 91–99.

32. Kwon Y, Ko H, Kim S, Kim M. The effect of cathepsin K inhibitor surface treatment on delayed tootg replantation in dogs. Dent Traumatol. 2018; 34(3): 201–7.

33. Andreasen JO, Borum MK, Jacobsen HE, Andreasen FM. Replantation of 400 avulsed permanent incisors. 4. Factors related to periodontal ligament healing. Endod Dent Traumatol. 1995;11: 76–89.

34. Miranda ACE, Habitante SM, Candelária LFA. Review of certain factors that influence in the success of the dental reimplant. Rev Biociênc. 2000; 6(1): 35–39.

35. Hinckfuss SE, Messer LB. An evidence-based assessment of the clinical guidelines for replanted avulsed teeth. Part II: Prescription of systemic antibiotics. Dent Traumatol. 2009; 25(2):158–64.

36. Guedes OA, Borges ÁH, Bandeca MC, de Araújo Estrela CR, de Alencar AG, et al. Analysis of 261 avulsed permanent teeth of patients treated in a dental urgency service. J Dent Res Rev. 2015; 2: 25–9.

37. Tsilingaridis G, Malmgren B, Skutberg C, Malmgren O. The effect of topical treatment with doxycycline compared to saline on avulsed permanent teeth. A retrospective case-control study. Dent Traumatol. 2015; 31(3): 171–6.

38. Kang W, Liang Q, Du L, Shang L, Wang T, Ge S. Sequential application of bFGF and BMP-2 facilitates osteogenic differentiation of human periodontal ligament stem cells. J Periodontal Res. 2019;54(4):424–34.

39. Santinoni CD, Oliveira HF, Batista VE, Lemos CA, Verri FR. Influence of low-level laser therapy on the healing of human bone maxillofacial defect: A systematic review. J Photochem Photobiol B. 2017; 169:83–9.

40. Acar AH, Yolcu U, Altindis S, Gul M, Alan H, Malkoc S. Bone regeneration by low-level laser therapy and low-intensity pulsed ultrasound therapy in the rabbit calvarium. Arch Oral Biol. 2016; 61:60–5.

41. Freitas NR, Guerrini LB, Esper LA, Sbrana MC, Dalben GDS, Soares S, Almeida ALPF. Evaluation of photobiomodulation therapy associated with guided bone regeneration in critical size defects. In vivo study. J Appl Oral Sci. 2018; 26: e20170244.

